# Protein-RNA condensation kinetics via filamentous nanoclusters

**DOI:** 10.1101/2024.11.05.622047

**Authors:** Ramon Peralta-Martinez, Araceli Visentin, Mariano Salgueiro, Silvina Borkosky, Mariana Araujo Ajalla Aleixo, Rodrigo Villares Portugal, Ignacio Enrique Sanchez, Gonzalo Prat-Gay

## Abstract

Protein-RNA phase separation is at the center of membraneless biomolecular condensates governing cell physiology and pathology. Using an archetypical viral protein-RNA condensation model, we determined the sequence of events that starts with sub-second formation of a protomer with two RNAs per protein dimer. Association of additional RNA molecules to weaker secondary binding sites in this protomer kickstarts crystallization-like assembly of a molecular condensate. Primary nucleation is faster than the sum of secondary nucleation and growth, which is a multistep process. Protein-RNA nuclei grow over hundreds of seconds into filaments and subsequently into nanoclusters with circa 600 nm diameter. Cryoelectron microscopy reveals an internal structure formed by incoming layers of protein-RNA filaments made of ribonucleoprotein oligomers, reminiscent of genome packing of a nucleocapsid. These nanoclusters progress to liquid condensate droplets that undergo further partial coalescence to yield typical hydrogel-like protein-RNA coacervates that may represent the scaffold of large viral factory condensates in infected cells. Our integrated experimental kinetic investigation exposes rate limiting steps and structures along a key biological multistep pathway present across life kingdoms.

## INTRODUCTION

Dynamic biochemical compartmentalization within cells is achieved by the formation of biomolecular condensates, based on liquid-liquid phase separation (LLPS) principles from polymer chemistry, and tightly associated with the concept of membraneless organelles^1,2^. Examples of biomolecular condensates relate to most aspects of biology both in physiology and pathology ^1,3–5^ Since the early discovery of the liquid nature of P granules^6^, protein-RNA condensation is at the center of most of these sub-cellular entities^7,8^, many associated to gene function^9^, stress and immunity^10–14^, and often associated with protein aggregopathies^15^. A wealth of reports described how it is modulated by RNA sequence, length, reentrant behavior, impacting on size, shape, viscosity, surface tension and composition^16–22^.

Viral replication sites, also known as viral factories, viral inclusion bodies, or viroplasms, were shown to be liquid in nature in rabies and respiratory syncytial viruses^23,24^, and since then, biomolecular condensation of viral factories appeared as a common theme among most viruses investigated to date, including SARS-COV2^25–31^. The nucleocapsid protein from SARS-CoV2 (N) was shown to be a main driver for liquid-liquid phase separation together with RNA^32–35^. Since the initial studies during pandemics, a vast number of reports on N condensation produced a wealth of information thoroughly reviewed and systematically compared by *Cascarina et al*. ^36^. Conclusions of this study are a fair summary of the current knowledge on the system : i) a role of N in virion assembly, genome packing and polymerase recruiting, ii) condensation is RNA dependent, electrostatically driven with little sequence specificity, iii) RNA concentration, length, base composition, and structure, can alter material properties and produce reentrant behavior, iv) although it bears an ideal LLPS prone architecture including RNA binding, dimerization, and substantial intrinsic disorder, there is no consensus on the participation of the different domains in condensation, v) proteolysis, mutation, and post-translational modification can influence N protein PS behavior *in vitro* and *in vivo*, vi) condensate morphology, size and material properties are highly condition sensitive and variable in different reports. The proposed and likely most relevant functions for this strong tendency of N to undergo heterotypic LLPS with RNA are recruitment of N protein to stress granules, selective condensation of the viral genome (gRNA), regulation of host-cell innate immune pathways, transcription modulation, recruitment of the viral RNA dependent RNA polymerase (RdRp) to viral replication centers, and anchoring of N-gRNA condensates at the endoplasmic reticulum-Golgi membrane prior to virus budding^36^.

Genome condensation can be linked both to replication by the polymerase and accessory proteins but also to the formation of a nucleocapsid that must pack into a virion. In single-stranded RNA virus assembly the genome packaging and capsid assembly are tightly linked and are considered to take place concomitantly^37^. This process is thought to involve weak protein-protein and non-specific electrostatic genome-protein interactions, including packing initiation signals in some cases, and participation of RNA secondary structure. A recent model paradigm involves the action of multiple cooperative contacts between nucleocapsid protein and RNA genome^37^. Some of these concepts are shared in protein-RNA condensation, and suggest a link between the two processes. A recent quantitative investigation of self-assembly kinetics of a viral capsid around its genome, revealed details on primary protein binding to the genome, via a nucleation and growth mechanism^38^.

In this work, we focus on N-RNA as a minimal model for investigating the kinetic mechanism, i.e. the sequence of events and species involved in a protein-RNA condensation reaction, which impacts in RNA virus gene function and assembly. We uncover the sequence of events that involve nucleation and a combination of first and second-order reactions, applying established models of self-assembly. We were able to obtain snapshots of intermediate species that can be characterized as self-limited and monodisperse nanuclustered condensates, with intriguing filament-like composition, which subsequently evolve into partially coalescing hydrogel-like coacervates. We discuss our findings in the light of fundamental protein-RNA condensation mechanisms and viral genome condensation and virion packing.

## RESULTS

### RNA binding and oligomerization equilibria preceding condensation of N

A myriad of reports with high variations in experimental conditions yielded different sometimes conflicting results, which suggest a high sensitivity of the system to the conditions^36^. In addition, to obtain a homogeneous and pure protein, free of any trace of RNA or conformational heterogeneities, we performed a stringent purification method involving two unfolding-refolding steps (see materials and methods). Two particularly sensitive parameters that impact on LLPS are the oligomeric state of the protein and RNA binding. The oligomeric state in our selected buffer conditions was evaluated by performing multi-angle light scattering size exclusion chromatography (SEC-MALS). The protein is present predominantly as a dimer (N_D_) (Figure 1A). However, variations in protein concentration indicate that a monomer-dimer equilibrium is established with a slow exchange between the species that allow to observe them as individual peaks (Figure 1B). Although the characteristics of the SEC technique does not allow to obtain an accurate *K*_D_, the ratio of both species shows a dissociation transition occurring with a midpoint around 0.5 μM of the injected sample (Fig. 1C). Considering a 20-fold dilution in the column, this would correspond to a *K*_D_ around 25 nM, in agreement with a previous report^39^.

**Figure 1.**
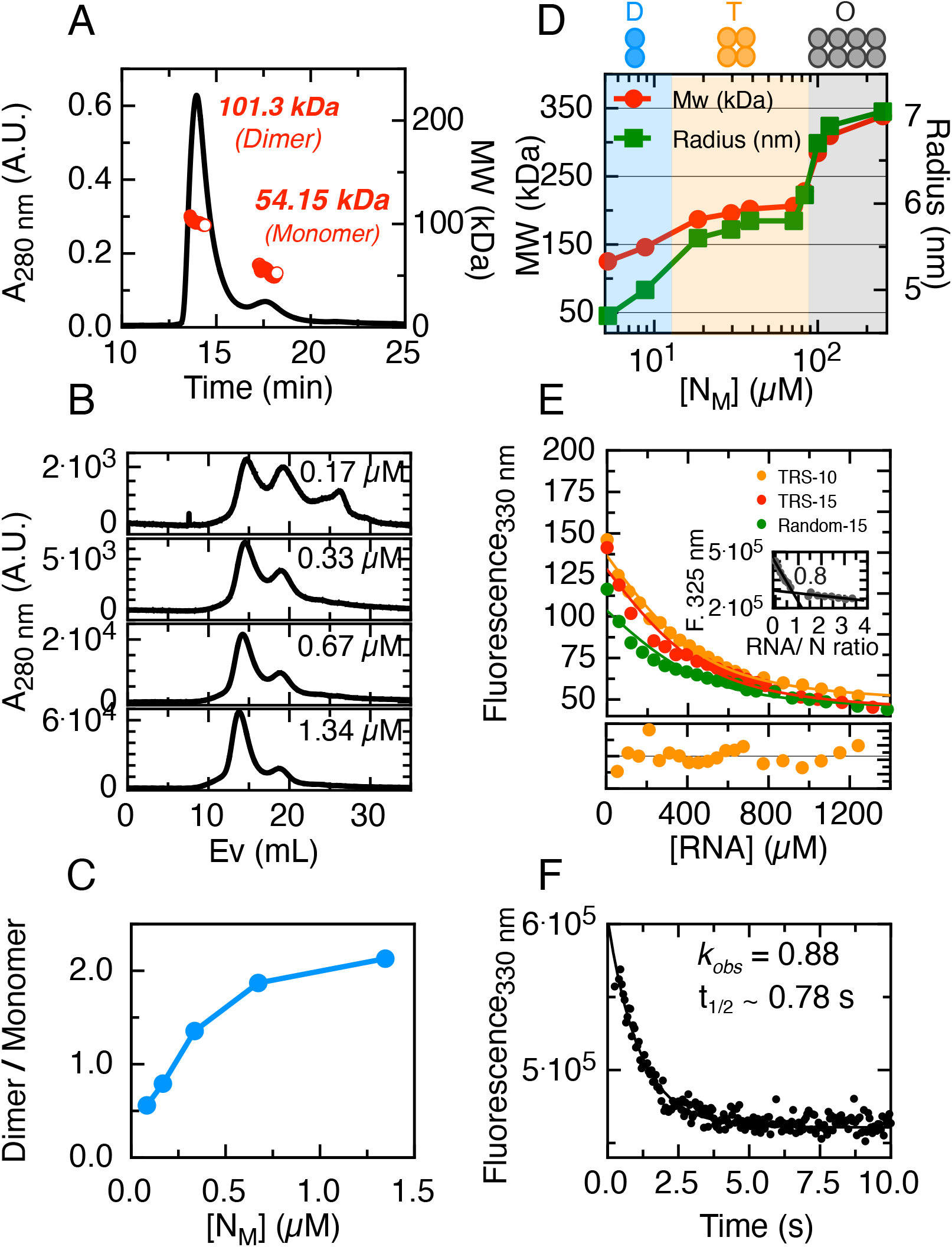
N protein oligomeric state is concentration dependant and it s binding to RNA is sequence inespecific. All protein concentrations used in this figure refer to monomer concentration [NM]. **(A)** Through SEC-MALS experiment the molecular weight of each of the peaks seen could be determined. UV 280 nm detection in black and molecular weight of each species detected in red. **(B)** Superdex 200 size exclusion chromatography (SEC) of the protein at different concentrations. **(C)** The ratio of Dimer/Monomer quantified in B was plotted for each protein final concentration. **(D)** Dynamic Light Scattering (DLS) experiments allowed us to study the oligomeric state of the protein for higher concentrations than those allowed in SEC and SEC-MALS. For each protein concentration both the molecular weight (green) and the radius (red) was measured. **(E)** Equilibrium binding assays where done for TRS-10, TRS-15 and Random-15 RNA s. Each experiment was done with a fixed protein concentration (500 nM) and tryptophan fluorescence was measured after the addition of increasing RNA concentrations. **(F)** Fluorescence trace of 0.4 μM protein after the addition of 50 nM RNA show a rapid binding kinetics, where the process reaches the equilibrium after five seconds.

In any case, is therefore safe to assume that in our experimental conditions the protein will not populate monomeric species (N_M_) above 2 μM. Dilution experiments using dynamic light scattering (DLS) allowed us to observe two consecutive transitions at higher protein concentrations (Figure 1D). At 5 μM the dimer is predominant, from 20 to 90 μM, the predominant species is a tetramer, and above 100 μM, an octameric species is formed.

RNA binding is also a fundamental aspect for the condensation mechanism, and therefore we tackled a quantitative assessment. For this, we followed the intrinsic fluorescence changes of the tryptophan signal upon the addition of RNA, which yields a substantial quenching. Three different RNAs were used: (i) TRS-10 (UCUCUAAACG), a 10-mer sequence previously described as relevant for coronaviruses life cycle^40^; (ii) TRS-15 (UUCUCUAAACGAACU), which extends five additional nucleotides, and (iii) Random-15 (AGUUGAGUUGAGUUG), a 15-mer RNA formed by a repeat of a 5-mer random sequence (AGUUG). We determined a binding stoichiometry of 1:1 N_M_:RNA from a titration/saturation experiment under this concentration regime (Figure 1E, inset). Under N-RNA dissociation conditions, we determined the affinity for the three different RNAs using a single-site binding model and observed no significant differences in affinity (Figure 1E), suggesting that the protein does not display high sequence specificity in short RNA sequences (10-15 mer). Additionally, to gain insight into the binding kinetics, we determined an observed rate of the RNA binding at 0.4 μM N_M_ and 50 nM of TRS-10 RNA to be 0.88 s^-1^, corresponding to a t_1/2_ of 0.78 s, indicating the primary is completed in under 5 seconds (Figure 1F).

### Modulation N condensation by short versus structured RNA models

As the oligomeric state of the protein depends on concentration, we decided to refer to monomeric concentrations in this section. The tendency for homotypic LLPS of N was evaluated by a phase diagram composed of protein and crowder concentration, where little if any droplets can be observed in the absence of crowder even at high concentration where the tetrameric species is expected (47 μM) (Figure S1A). In the presence of PEG, we observe a transition from small bead like sticky droplets to regular spherical droplets suggestive of different physicochemical properties between 1.9 and 3.8 % crowder (Figure 2A and 2B).

**Figure 2:**
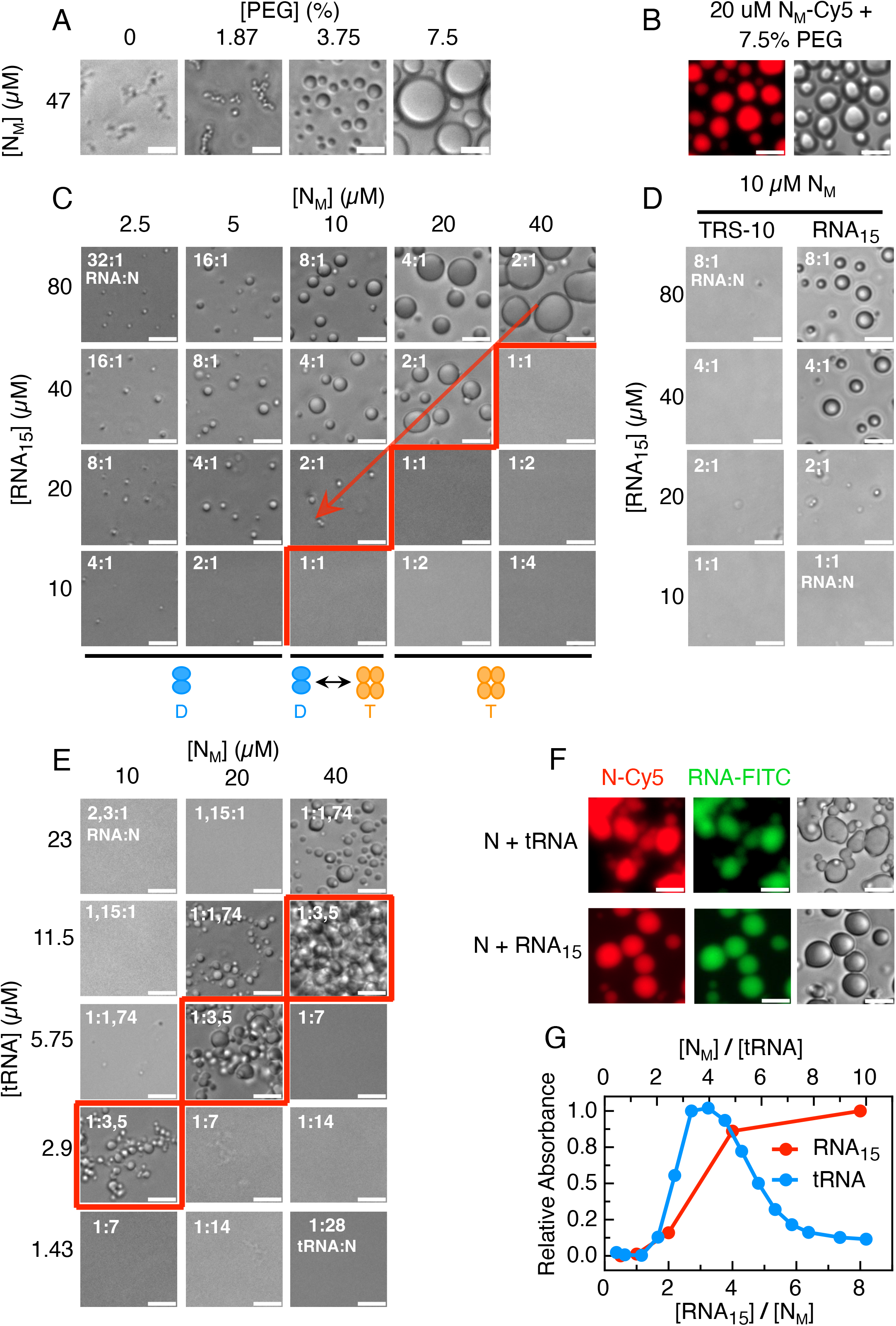
N forms homotypic and heterotypic condensates in different stoichiometric ranges. **(A)** All protein concentrations used in this figure refer to monomer concentration [NM]. Light microscopy images of homotypic condensates formed by N in presence of crowding agent PEG 4000. **(B)** Fluorescence microscopy images of N-Cy5 forming homotypic condensates. **(C)** Light microscopy images showing heterotypic condensates formed in different concentrations of protein and RNA15. **(D)** When using a RNA10 no condensates are formed. **(E)** Heterotypic condensates formed with total yeast tRNA. **(F)** Co-localisation of protein and RNA seen in through fluorescence microscopy of N-Cy5 with fluorescent labeled RNA (tRNA or RNA15). **(G)** Light scattering at 370 nm show the different stoichiometric windows where heterotypic condensates are formed, both for N-RNA15 (red) and N-tRNA (blue). In all images scale bars are 10 μm.

Next, we analyzed the formation and modulation of heterotypic N:RNA LLPS by performing phase diagrams varying the concentration of both components. The 15mer RNA (with no difference in binding affinity to TRS15, Figure 1E) form coalescent droplets increasing size towards high protein and RNA concentration as expected (Figure 2C). Noticeably, there is a sharp transition boundary that shows a condensation onset at 2:1 RNA:N (Figure 2C, red boundary) and not 1:1 RNA:N stoichiometric binding ratio (Figure 1E), not even at high concentrations (Figure 2C). Below the red boundary no LLPS is observed in excess N, and the opposite applies to excess RNA above the red boundary. Condensates at 2 RNA_15_: 1 N ratio disappear upon dilution (Figure 2C diagonal arrow down). The propensity of N to condensate with RNA_10_ is negligible compared to the RNA_15_ under a similar concentration regime (Figure 2D), and requires lowering the ionic strength (Figure S1B).

N protein wraps and condenses the 30kb genome of the virus and bind to RNA transcripts, implying interaction with longer and heterogeneous sequences. In order to test a simple model for heterogeneous variable sequences and structure encountered in the cellular environment, we make use of total purified yeast t-RNA, with an average length of 70 bases. The phase diagram shows that condensation has an optimal stoichiometric window of 1 tRNA : 3.5 N (Figure 2E, red boundaries), and transition to one-phase above and below that ratio. The difference between RNA_15_ and tRNA is clearly observed when we monitor turbidity along the RNA : protein ratio range, showing a finely tuned reentrant phase for the tRNA but not for the 15mer (Figure 2C, 2E, and 2G). A noticeable observation is that RNA_15_-N condensates are regular coalesced droplets, while the tRNA-condensates have a more rigid appearance that tend to stick with only partial coalescence at higher concentration, typical of hydrogel-like coacervates. We confirmed that both protein and RNA are present in the droplets of the RNA_15_ as well as tRNA (Figure 2F), but we choose not to use chemically modified N to avoid or minimize possible effects of the fluorophore labelling.

While the tRNA-N condensates are highly sensitive to an increase in the ionic strength (Figure 4A left) as expected for a charge coacervate of oppositely charged polymers, the crowder enhanced homotypic condensate is increased at high salt (Figure 4A right), strongly suggesting that condensation leads on average to formation of unfavorable electrostatic interactions relative to N in solution. Conversely, the heterotypic condensate is less stable at high salt concentrations. This suggests that condensation of N and tRNA, which have charges of opposite signs, leads on average to formation of favorable electrostatic interactions relative to the two molecules in solution. Overall, these set of experiments allow us to define controlled solution conditions and macromolecular ratios, based on the different phase diagrams observed, for tackling a quantitative mechanistic investigation

### Kinetic mechanism of N-RNA condensation

Comprehensive understanding of a condensation mechanism involves the dissection of the initial, intermediate and endpoint species present, their rates of conversion, and the sequence of events. To this end, we investigated the evolution of the turbidity with time measured as scattered absorbance at 370 nm light, indicative of condensation (Figure 2G). We triggered the reaction by adding RNA_15_ (Figure 3A) or tRNA (Figure 3B) to pre-incubated N at different concentrations, while keeping a RNA:N ratio (2:1 and 1:3.5 respectively, Figures 2C and 2E). We monitored the change in the absorbance signal over hundreds to thousands of seconds (Figures 3A and 3B), and no evidence of fast burst-phase events in terms of condensation in either case, as no significant changes in absorbance can be observed within the experimental deadtime (ca. 15 seconds). The kinetic traces of both condensation processes can be described empirically using a sum of two exponential reactions (Figure S2A for RNA_15_ and S2B for tRNA). The fitted amplitudes of the fast and slow processes phases are close to 50% each. The empirical rate constants in the case of N-RNA_15_ condensation are 0.0085 ± 0.0001 s^-1^ (*k*_obs1_) and 0.00084 ± 0.00001 s^-1^ (*k*_obs2_) at 1 μM N_D_ (4 μM RNA-15). Rates of 0.0081 ± 0.0002 s^-1^ (*k*_obs1_) and 0.0021 ± 0.0001 s^-1^ (*k*_obs2_) were determined for tRNA condensation, at 1 μM N_D_ (0.6 μM tRNA). Thus, although no significant differences are observed for *k*_obs1_, tRNA condensation showed a *k*_obs2_ almost three-fold higher compared to RNA_15_ condensation. Moreover, the total amplitude of both processes decreases at low protein concentrations and display a similar condensation onset at ca. 0.7 μM N-RNA (Figure S2C). No lag phase was observed for N condensation under these conditions (Figures 3A and 3B). However, a lag phase could take place within our experimental deadtime, depending on the conditions and the values of the rate constants.

**Figure 3:**
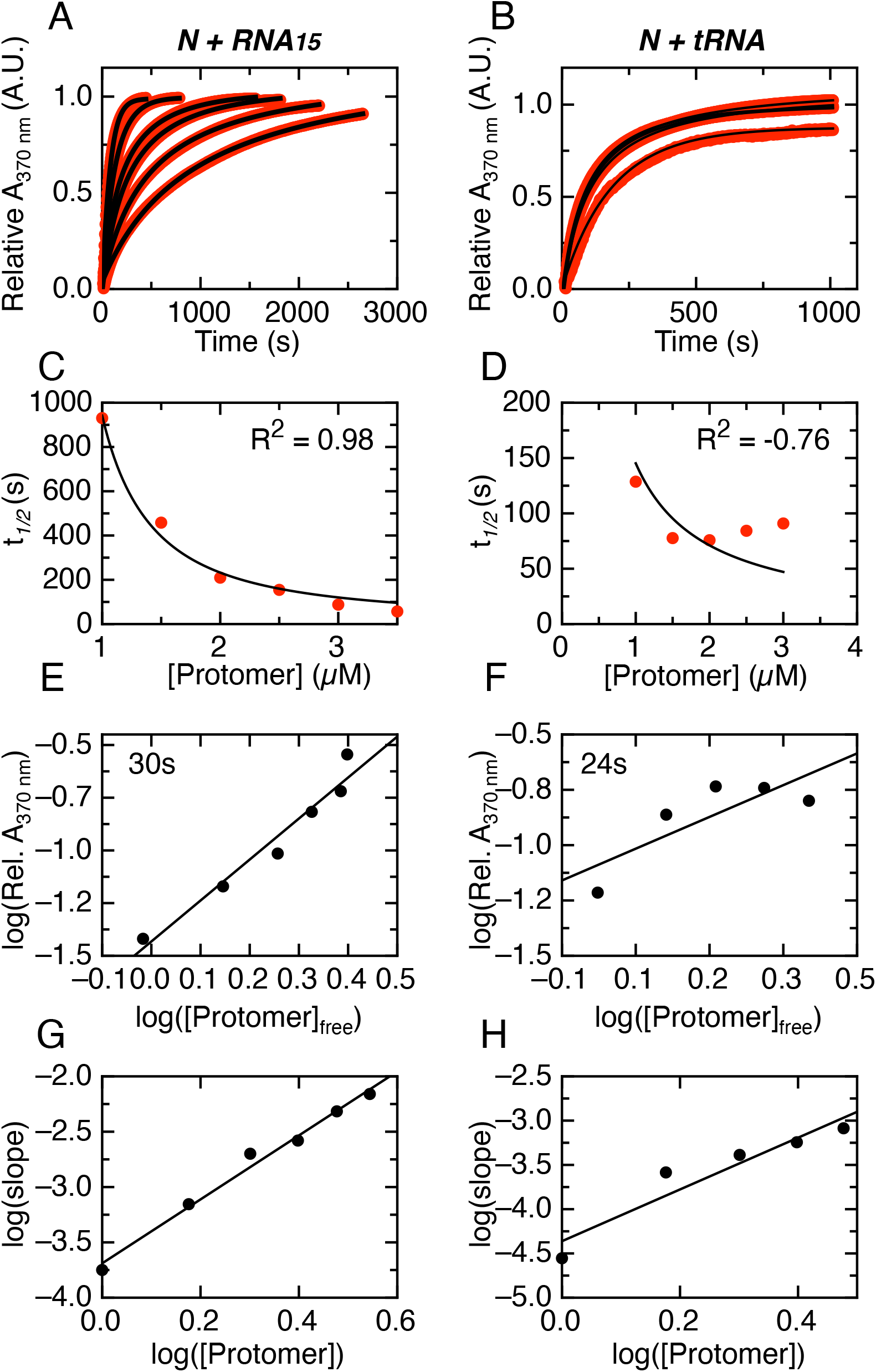
The nucleus size and concentration dependance of heterotypic N condensation is RNA dependent: As model analysis depends on protein oligomeric state, all protein concentrations used in this figure refer to protomer concentration. Normalized 370 nm light scattering traces of N-RNA15 **(A)** and tRNA **(B)** condensation for varying concentrations of a fixed N:RNA ratio. In black, hyperbolic fittings from NAGKPin software are shown. **(C-D)** Crystallization-like assembly model fitted to the *t*1/2 obtained from the fittings in A-B for N-RNA15 and tRNA respectively. **(E-F)** Zlotnick plots for determination of the nucleus size for N-RNA15 and tRNA respectively. The nucleus size is obtained from the slope of the linear fittings, being 1.94 ± 0.21 for N-RNA15 condensation and 0.96 ± 0.40 for tRNA condensation. **(G-H)** Zlotnick plots for determination of the elongation reaction order for N-RNA15 and tRNA respectively. The elongation order is obtained from the slope being 2.89 ± 0.17 for N-RNA15 condensation and 2.92 ± 0.55 for N-tRNA condensation.

To gain insight on the N-RNA condensation mechanism, we first applied the model by Martins and coworkers applied to the crystallization-like assembly originally developed to describe amyloid fibril formation but may be used to study any association process of multiple molecules ^41 42^. This model describes condensation by a general mechanism involving supersaturation-dependent nucleation and growth steps. The concentration dependence of condensation can be used to extract information of primary and secondary nucleation, growth and the presence of parallel pathways. The model considers the formation of a primary nucleus from molecules in the solution, growth of the primary nucleus and the formation of secondary nuclei on the surface of the primary nucleus. The analysis of experimental data quantifies the relative importance of the kinetic steps of primary nucleation, secondary nucleation and growth. We fitted the model to normalized mass-progress curves (Figure 4A and 4B, data in red, fitting in black) and to the scaling of *t*_1/2_ versus the initial N dimer concentration (Figures 3C and 3D). The model describes the concentration dependence of N-RNA_15_ condensation very well (R^2^ 0.987, Figures 3C), with a fitted value for the critical solubility (Cc) of 0.698 μM N_D_ in excellent agreement with that obtained from extrapolation of the total condensation amplitude (Figure S2C). The fitted autocatalytic rate (*k*a) is 0.0031 s^-1^, which corresponds to the sum of the rate constants for growth (*k*_+_) and secondary nucleation (*k*_2_). Mass-progress curves alone are unable to distinguish between these two processes, which requires a detailed analysis using multiple size-progress curves at different concentrations. The dimensionless nucleation rate (*k*_*b*_=*k*_*n*_/*k*_*a*_) is 0.999. This value is greater than 0.1, indicating that according to the model, the primary nucleation is faster relative to the sum of rate constants for growth and secondary nucleation. A global fit of the data yielded a r^2^ value lower than 0.95 (not shown), suggesting plausible parallel processes such as coalescence and/or off-pathway condensation. Overall, we find that N-RNA_15_ condensation can be described using the crystallisation-like assembly model, where primary nucleation is relatively fast and parallel processes are likely present.

On the other hand, the model describes the concentration dependence of N-tRNA condensation poorly, due to the process slowing down from 2 to 3 μM N_D_ (Figure 3D, R2 -0.76). The poor quality of the fit prevented us from extracting values for the autocatalytic rate (ka) or the dimensionless nucleation rate (*k*_*b*_=*k*_*n*_/*k*_*a*_). However, the v-shaped dependence of *t*_1/2_ does have relevant mechanistic information, since it suggests the dominance of secondary nucleation over nucleus growth in the the crystallisation-like assembly process and the additional presence of soluble and stable off-pathway aggregates (see Figure 6, panels E to F in *Silva et al*.^43^).

**Figure 4:**
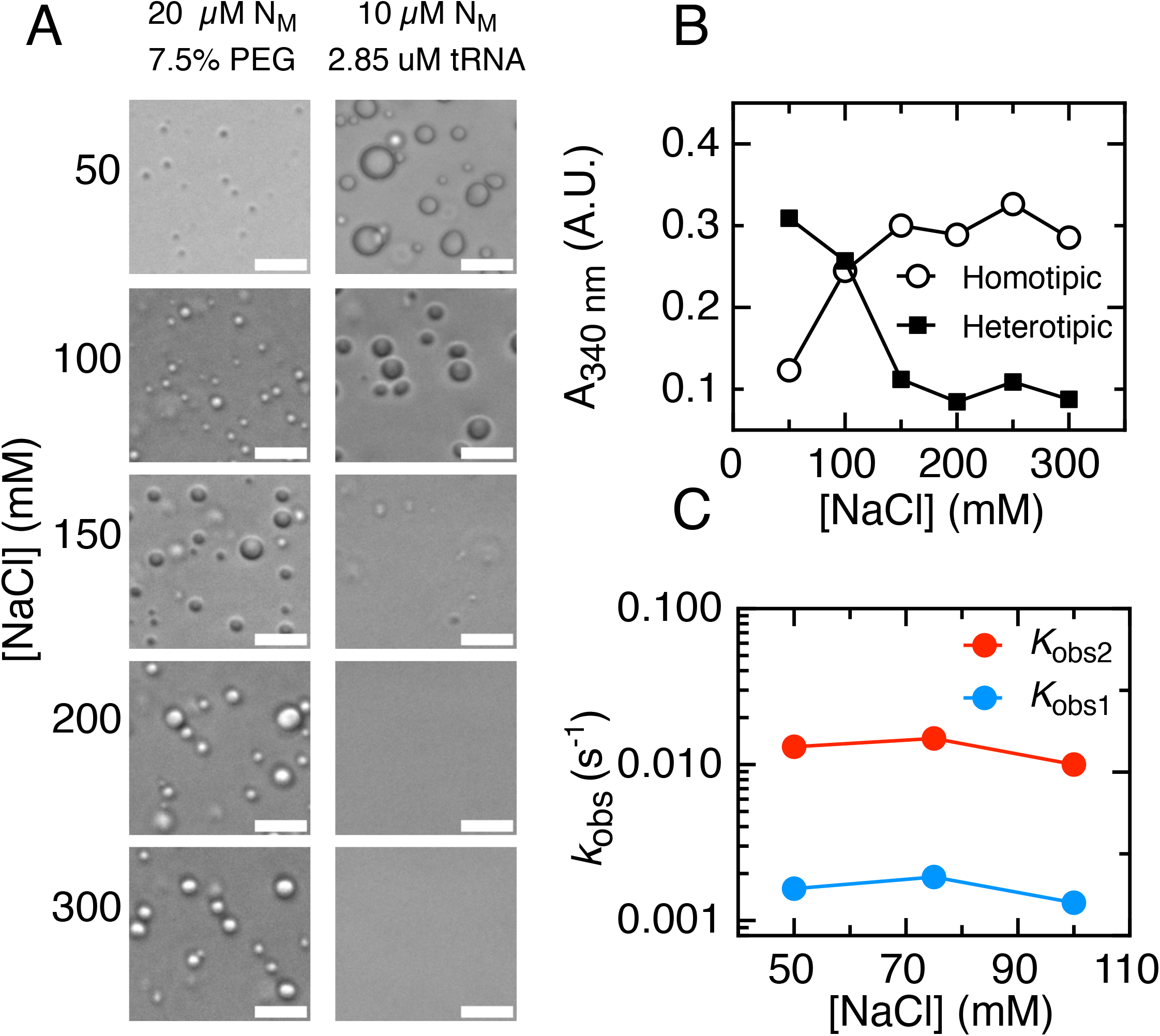
Effect of ionic strength on condensates. All protein concentrations used in this figure refer to monomer concentration [NM]. **(A)** Light microscopy images of homotypic (20 μM N + 7.5% PEG 4000) and heterotypic (10 μM N + 2.85 μM tRNA) condensates with increasing concentrations of NaCl. **(B)** 370 nm light scattering of homotypic (20 μM N + 7.5% PEG 4000) and heterotypic (10 μM N + 2.85 μM tRNA) condensates. **(C)** Observed rate constants of N-tRNA condensates for varying NaCl concentrations. These rates were obtained from the double exponential fitting of 370 nm scattering traces. In all images scale bars are 10 μm.

As a complementary approach, we made use of the model by Zlotnick and coworkers^44^ for the kinetically limited assembly of a molecular condensate. Although this model was originally developed to describe the assembly of viral capsids, it may be used to study other association processes. The kinetically limited assembly model describes condensation as a cascade of low-order association reactions, where a rate-limiting “nucleation” step is followed by faster elongation steps.

The concentration dependence of condensation can be used to extract information on both the nucleation and the elongation steps. The size of the nucleus can be calculated from the slope of a double logarithmic plot of the concentration of condensed versus free N-RNA at a fixed time point early in the reaction, typically at a time when the kinetics is well described by a straight line^44^. In the case of N-RNA_15_ and N-tRNA condensation, we calculated the concentration of condensed and free N-RNA in the linear range (see Experimental Procedures). Using this approach, we obtain a slope of 1.94 ± 0.21 for N-RNA_15_ condensation corresponding to a nucleus formed by N-RNA_15_ protomers (Figure 4E), each protomer corresponding to the stable Ndimer-RNA_15_ stoichiometric complex (Figure 1F, i.e.), referred to as the nucleation protomer. In the case of N-RNA_15_ reaction, the nucleus is composed of two protomers.

In the case of N-tRNA condensation, a slope of 0.96 ± 0.40, indicates a nucleus formed by one N-tRNA dimer as the protomer (Figure 4F). We interpret that a slow early step in N-RNA_15_ condensation likely involves association of two N dimers after RNA binding, and a conformational rearrangement of the initial N dimer after tRNA binding for N-tRNA condensation.

Moreover, the reaction order of the faster elongation step can be determined from the slope of a double logarithmic plot of the initial rate in the reaction (typically at a time when the kinetics are well described as a straight line) versus the initial concentration of N dimer protomers. This analysis yields slopes of 2.89 ± 0.17 and 2.92 ± 0.55, for N-RNA_15_ and N-tRNA condensation, respectively (Figures 3G and 3H). This indicates that the reaction order of the elongation reactions for both N-RNA_15_ and N-tRNA condensation is close to three. There are two possible interpretations for this number. The first one is that the elongation step is a single-step, elementary reaction that involves simultaneous collision of three molecules protomers. We interpret this as improbable because ternary collisions with the right orientation and energy are rare and the resulting reaction would likely be slower than the competing processes. The second interpretation is that the elongation step in both N-RNA_15_ and N-tRNA condensation is a multi-step process that involves the incorporation of multiple N-RNA protomers to the growing nucleus. In summary, the kinetically limited assembly model allowed us to describe the nucleation and elongation steps for both N-RNA_15_ and N-tRNA condensation. The two processes show clear differences, and additional complexities to be characterized in future work.

To analyze the contribution of electrostatic interactions to the kinetics of N-tRNA condensation, we evaluated the effect of modifying the salt concentration on the observed rates from exponential analysis (Figure 4C). The absorbance amplitude of the reaction decreased with increasing sodium chloride concentrations (Figure 4B), as expected from the salt dependence of the condensation droplets observed under the microscope (Figure 4A). The two fitted rate constants remain approximately constant upon going from 50 to 100 mM sodium chloride. Since sodium chloride does not have a net effect on the kinetics of N-tRNA condensation, we interpret that the electrostatic interactions stabilizing the condensate are not formed in the rate-limiting transition states along the pathway.

### Characterization of intermediate species along the kinetic condensation pathway

We next tackled the characterization of the intermediates involved along both N-RNA condensation pathways, by analyzing the sizes of the species involved using dynamic light scattering (DLS). We used a N concentration of 5 μM monomer, which allows for an accurate determination of sizes and using the same N:RNA ratio as in the kinetic experiments. Under these conditions, the hydrodynamic radius (*r*_h_) of the uncondensed N dimer is between 6.5 and 7.0 nm (Figure 5A and 5B, grey peaks. Upon addition of RNA_15_ or tRNA, we observe the formation of larger mono-disperse species within two minutes, with *r*_h_ of 120 and 261 nm, respectively (Figures 5A and 5B, 2 min red and blue). The hydrodynamic radius of these species increases to 320 nm for RNA_15_ and 836 nm tRNA after 30 minutes. For both reactions, we observe intermediate sized species of 22.8 ± 2.51 nm and 46.03 ± 5.38 nm in average for RNA_15_ and tRNA, respectively (Figures 5A, magenta peaks in the insets). Noticeably, the peaks are mono-disperse, indicative of discrete sizes, compatible with a nucleation-growth model. However, tRNA peaks show increased poly-dispersity as time increases. Finally, at the longer times we observe a poly-disperse species larger than 2 μm in size, above the limit of the instrument measurement (Figures 5A, orange peak in insets). We ascribe this species to droplet coalescence events, which become evident in microscopy when the droplets decant.

**Figure 5.**
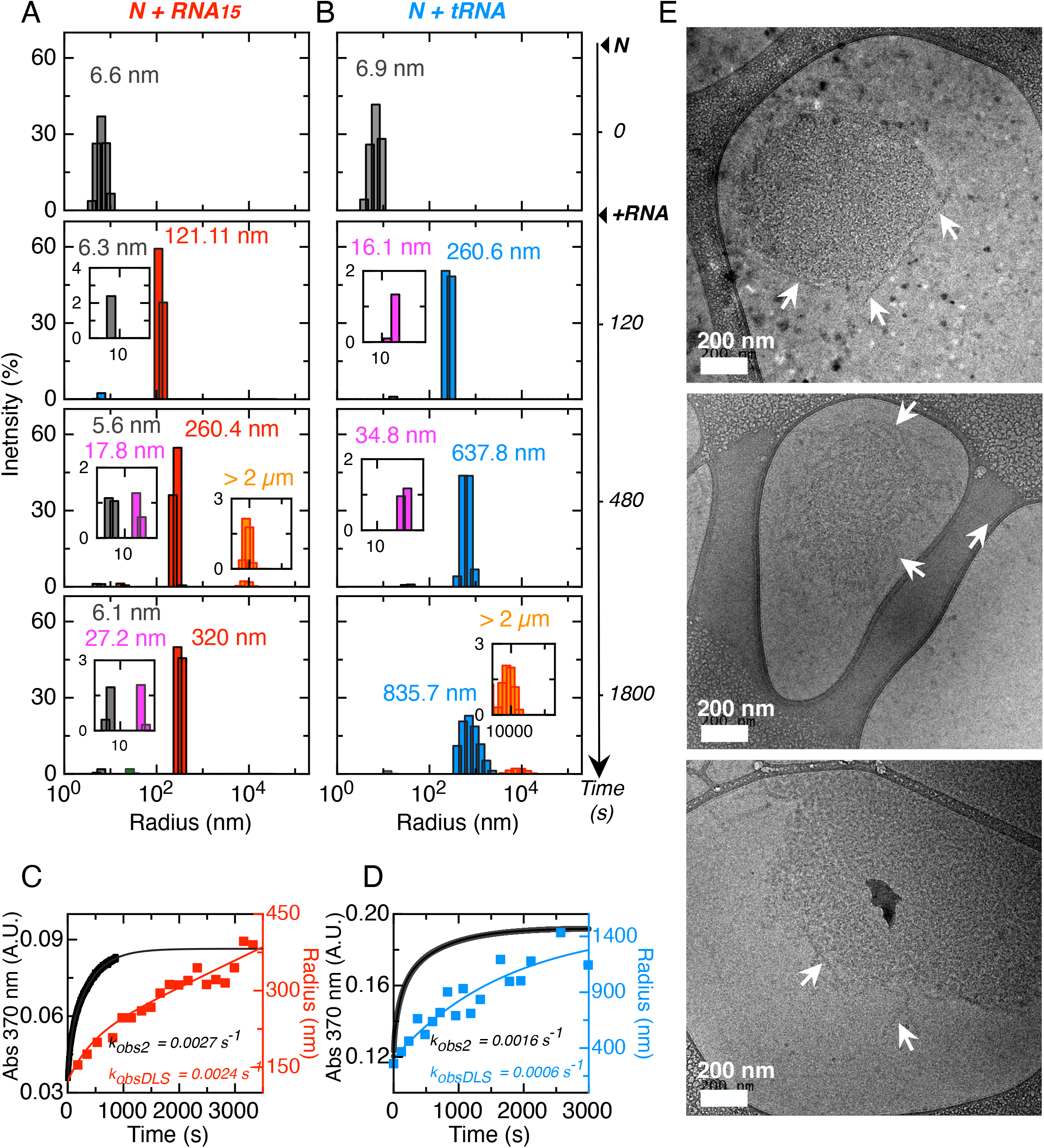
Dynamic Light Scattering histograms corresponding to different times of a RNA15 **(A)** and tRNA heterotypic condensation reaction **(B)** At 0 seconds the histogram of the protein only can be seen. Then, the correspondent RNA concentration was added in each case and several histograms where obtained (from 120 seconds onwards). **(C-D)** The radius of the population coloured in A as red (for RNA15) and blue (for tRNA) was plotted against time. **(E)** Cryo-EM images of 5 μM N + 20 μM RNA15 after 45 minutes of reaction. The size bars correspond to 200 nm. The white arrows indicate the filaments

The evolution of the radius of the major species was also followed in detail in 10 second intervals by DLS, with the data shown in Figures 5B and 5C, overlayed to mass-progress turbidity curves measured in the same conditions. The time evolution of *r*_h_ could be fitted to a single exponential function for both N-RNA_15_ condensation (*k*_*DLS*_ of 0.0024 ± 0.003 s^-1^) and N-tRNA condensation (0.0006 ± 0.00045 s^-1^). These values are in very good agreement with the rate constants for the slowest process in the absorbance traces (*k*_obs2_) under the same conditions, which are 0.0027 ± 0.0002 s^-1^ for N-RNA_15_ condensation and 0.0016 ± 0.00045 s^-1^ for N-tRNA condensation. This strongly suggest that the slowest process in the mass progress curves corresponds to the growth of condensates.

With the goal of obtaining a further layer of structural information about the condensation intermediates we tackled their analysis by Cryo Electron Microscopy (Cryo-EM). We triggered N-RNA_15_ condensation by adding 20 μM of RNA_15_ to 5 μM N, and flash-froze the reaction after 30 min, where the turbidity experiments reach a steady state (Figure 3A). Two dimensional Cryo-EM images show semi-regular spherical structures of sizes ranging from 300 to 700 nm (Figure 5D), in agreement with mono-disperse species found by DLS, considering the expected differences between techniques. Interestingly, close inspection of the images shows a marked internal arrangement in the shape of layered filaments as opposed to the homogeneous aspect one would expect from a liquid. These filaments are clearly in close contact and in a defined arrangement. Although their precise size and distribution would merit a more detailed investigation, their length oscillates between 180 and 300 nm, but the width is more homogeneous, with a value of 27 ± 3 nm (Figure 5D). Moreover, in all images inspected the edges of these structures show filaments shedding off the border reminiscent of a wool ball shape (Figure 5D, white arrows). We ascribe these to incoming loose filaments in the process of formation of these structures. Control images with either N or RNA alone show no evidence of any type of structure (Figure S3), suggesting that the filaments originate from N-RNA interactions. Thus, Cryo-EM images of N-RNA_15_ revealed filamentous structures which size is in excellent agreement with the mono-disperse species detected by DLS.

## DISCUSSION

In this work, we have dissected the kinetic condensation mechanism of N-RNA using mass-progress curves, size-progress curves, a crystallization-like assembly model, a kinetically limited assembly model, DLS, and cryoelectron microscopy. In our working conditions, N is a dimer and binds RNA with undifferentiable affinities (ca. 165 nM) between cognate specific and unrelated sequences. The difference in condensation and binding stoichiometry reflects the presence of secondary lower affinity RNA binding sites that operate as the protein concentration increases. Thus, primary RNA binding and condensation reactions are well separated by time and by concentration.

We made use of two different types of RNA to investigate the mechanism. The 15-mer RNA oligonucleotide yielded a fine stoichiometric dependance for condensation to occur showing a threshold at 2RNA_15_:1N ratio, with no reentrant behavior. The heterogeneous and structured ca. 70 base tRNA also showed a stoichiometrically controlled condensation but displayed a strong reentrant behavior that supports the idea that the modulated assembly-dissolution of tRNA condensates represents a better model for the in-cell scenario. The lack of sequence discrimination capacity under condensation concentrations, suggests that the protein would not discriminate genomic from transcript RNA.

By combining DLS and cryoelectron microscopy we were able to characterize the nature of the condensation intermediates. Both RNA model reaction pathways populate a major intermediate species with remarkable mono-dispersity within 30 minutes of reaction, particularly for the chemically homogeneous RNA_15_. On the other hand, Cryo-EM images revealed roughly rounded shaped particles of sizes compatible with those from DLS (300 to 600 nm diameter), composed of a surprising network of layered filaments. These filaments vary in length but show homogeneous widths (ca. 27 nm), however, further structural analysis should be tackled to reveal structural details. We propose that the frayed edges observed (Figure 5E) correspond to incoming filaments on the way to the early particle growth, in agreement with our experimentally based kinetic model supported by DLS. We know that the subsequent event down the pathway is the coalescence into droplet coacervates, which require the fibrillar intermediates to be assembling rather than disassembling. The layered filamentous organization of these particles as opposed to a homogenous density phase^45^ suggests a non-liquid nature, which makes us consider these structures as nanoclusters. Altogether, N-RNA_15_ condensation proceeds through a filamentous nanoclsuter intermediate, where the multiple oligomeric species in equilibrium we and others^39,46^ observed for N across the concentration range and stabilized by RNA are the building blocks for the filaments.

We combined our results in a full mechanistic model described in Figure 6. N-RNA_15_ condensation requires fast primary binding of RNA, which takes place in the sub-second time window, typical of an electrostatically driven collision, largely preceding condensation. At concentrations under saturation, we expect the main binding event to be that of the RBD. As concentration increases over saturation, we expect additional RNA binding events to occur at low-affinity sites present in other regions of the protein^47^. The saturation of these sites triggers the formation of a nucleus formed by two protomers. This resembles what was previously reported^39^, where in excess of RNA a tetramer is formed with a 2:1 binding stoichiometry, which correlates with the 2:1 condensation ratio we observed (Figure 2C). This nucleation process is fast relative to the processes of secondary nucleation and growth. The relationship between the nucleus and the native tetramer observed at higher concentrations remains to be elucidated. The transition from the nucleus to the larger condensates observed in light-microscopy goes through a multi-step elongation reaction, involving the formation of a filamentous intermediate which further associates into a nanocluster. The further growth of these nanoclusters likely involves parallel processes such as coalescence and/or off-pathway condensation.

An increase in ionic strength destabilizes the condensate but does not affect the condensation kinetics, suggesting that the overall favorable interactions between the negatively charged RNA and the positively charged N consolidate only in the final condensate. Moreover, the tendency of the protein to form homotypic condensates in the presence of crowder and the insensitivity of these to ionic strength reveals that hydrophobic protein-protein self-interaction likely plays a role in heterotypic condensation with RNA. Taking this together, considering also the inability of the protein to condensate with RNA_10_, N-RNA condensation occurs through a combination of homotypic and heterotypic interactions as previously reported ^48^.

**Figure 6:**
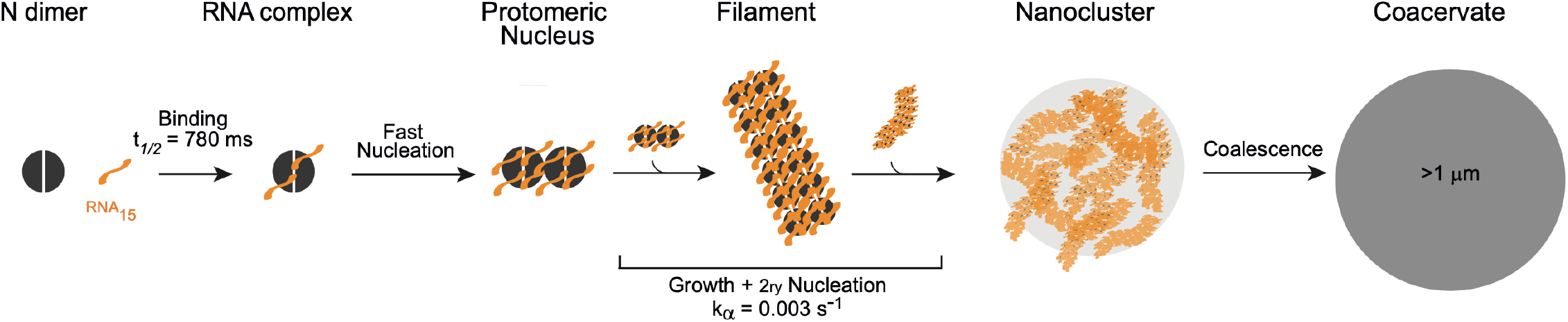
Mechanistic model of RNA15 condensation. After sub-second binding events, fast nucleation occurs relative to secondary nucleation and growth. This nucleus is formed by two protomers. Nucleus growth occurs through a filamentous intermediate, which further associates to form a filamentous nanocluster. This nanocluster grows by addition of filaments until adopts liquid properties which allows it to coalesce forming liquid condensates as seen in light microscopy images.

Interestingly, the difference in the concentration dependence of the condensation kinetics of both RNA studied reveals that the molecular details of the process, including nucleus size, rate constants, and the presence of off-pathway intermediates are dependent on the identity of the condensing RNA molecule. Thus, on one hand the occurrence of condensation shows little sequence specificity and seems to be a robust feature of the virus life cycle. On the other hand, the exact way in which it takes place may be perturbed by changes in the RNA, arising from natural mutations, cellular state, or other external factors.

The monodispersity of the nanoclusters observed by DLS supports a nucleation mechanism where particles grow from individual nuclei to reach a size plateau. This also suggests that these structures are self-limited in size. These structures resemble the internal organization in virion nucleocapsids^49^ and the self-limitation further supports that the reaction may represent the condensation of a packed genome in the path to virion formation. These nanoclusters are significantly larger than the virion, 300 to 700 nm in diameter compared to 50 nm. However, we highlight the capacity of self-limitation and the presence of internal capsid-like structure. We speculate that large liquid-like condensates represent the viral replication factories as large reaction vessels in a different liquid phase, where most viral RNA synthesis takes place (ref), whilst the self-limited layered nanoclusters may represent the type of structural organization in a virion proto-nucleocapsid. These events could be sequential, i.e., the nanoclusters can either evolve to large liquid condensates or be an endpoint in nucleocapsid formation. The fate of each pathway may be dictated by viral and host factors, or whether the composing RNA is either transcript or genomic. Since there is little if any RNA sequence specificity condensation, there must be other distinctive features in both RNAs, including expression timing and modification of their relative levels.

In this work, we uncovered an elementary mechanistic pathway in a protein-RNA condensation reaction, impacting the myriad of biomolecular condensates linked to cell physiology and pathology, particularly in early events and the definition of sequential steps. In addition, our findings have direct impact in SARS-CoV-2 N protein and N-RNA condensation in RNA viruses in general. Understanding biological biomolecular condensation at the molecular and physicochemical level, requires mapping experimental kinetic pathways with starting, intermediate and final species, their energy barriers, and the thermodynamics involved. The ability to dissect and reproduce a robust protein RNA condensation reaction contributes to the ability to design and tune condensates with further relevance for nanotechnology and synthetic biology.

## MATERIALS AND METHODS

### Protein expression and purification

The sequence expressing SARS-CoV-2 N protein was obtained from Genscript inside Pet-28 plasmid. N protein vectors were transformed into *E. coli* C41 for expression. Freshly transformed cells were grown at 37°C in LB-Kanamycin until OD 0.8. Protein expression was the induced with 1 mM IPTG for 12 h. Harvested cells were resuspended in Lysis Buffer (Tris-HCl 50 mM pH 8, NaCl 1M, PMSF 1mM) and sonicated. 6M Urea was added to the soluble fraction, left overnight at 25oC and loaded on to a Ni2+ affinity column in Buffer Sodium Phosphate 50 mM pH 8, NaCl 0.4M, Urea 6M. Protein was then eluted with Buffer Sodium Phosphate 50 mM pH 8, NaCl 0.4M, Urea 6M, 400 mM imidazole and then dialysed in Buffer Sodium Phosphate 50 mM pH 8, NaCl 0.4M. For His-Tag cleavage and removal, protein was incubated overnight with 1:500 TEV : N at 25°C. Then 6M Urea was added to the solution, left overnight at 25°C and loaded on to a Ni2+ affinity column in Buffer Sodium Phosphate 50 mM pH 8, NaCl 0.4M, Urea 6M. The cleaved protein was collected in the flow-through and then dialysed in Sodium-Phosphate 25 mM pH 8, NaCl 0.2M. Protein was concentrated using Millipore Centrifugal Amicon filters and stored at -80°C until use.

### Size Exclusion Chromatography and SEC-MALS

Buffer Phosphate 25 mM pH 8, NaCl 0,4M was used for both SEC-FPLC and SEC-MALS experiments. For SEC-FPLC, Superdex-200 column was used in Shimadzu SPD-10A equipment. For SEC-MALS, Wyatt Mini DAWN + Optilab equipment coupled to a Jasco UV-4075 HPLC was used. For this a Wyatt 5-1.250 kDa - 500 Å pore size was used.

### Equilibrium binding assays

For determination of the apparent dissociation constant, the concentration of N protein was fixed in 500 nm and tryptophan fluorescence signal was traced as increasing concentrations of RNA were added. Sample was excited with 290 nm and emission in 325 nm was detected. The excitation slit was set in 5 nm and emission slit in 8 nm. Blank was subtracted and the data was fitted to a reversible one-site model using Pro-Fit software (QuantumSoft).

### Light microscopy

For microscopy imaging 96-well non-binding bottom Greiner plates were used. HEPES 20 mM pH 7.5, NaCl 100 mM was used (unless otherwise indicated). Reactions were triggered after the addition of RNA and incubated for 30 min at room temperature before imaging. The images were acquired using an Axio Observer 3 inverted microscope with a 40x/0,750.3 M27 objective and a Colibri 5 LED illumination system if needed. Images were processed using Fiji (A distribution package of ImageJ software, USA).

### Phase separation kinetic assays

Scattering measurements were done in Jasco V-750 Spectrophotometer setting A370 nm. Reactions were done in HEPES 20 mM pH 7.5, NaCl 100 mM 25°C. Data was further analyzed. For mechanistic analysis, turbidity kinetic traces were fitted using a two-exponential function plus a drift, using the following equation 2 *y* = *A1* exp(*k*_obs1_ *t*) - *A2 exp*(*k*_obs2_ *t*))+m *t*, where A_*i*_ is the amplitude of the phase *i* and *k*_obs*i*_ its empirical rate constant.

### Dynamic Light Scattering measurements

DLS measurements were carried out on DynaPro NanoStar II DLS device from (Wyatt Technology). Phase separation measurements were performed HEPES 20 mM pH 7.5, NaCl 100 mM. For this, 5 μM N was first measured to obtain the protein only measurement. Then either 20 μM of RNA_15_ or 1.4 μM tRNA were added and its kinetics were followed by doing 30 measurements every 12 s, each composed of 5 acquisitions with 20 s acquisition time. The temperature was maintained at 25°C by Peltier control system. Results were processed employing the software package included in the equipment. DLS trace was fitted using a single-exponential function using the following equation *y* = *A* exp(*k*_obs_ *t*) + m *t*.

### Cryo-EM

The condensation process was trigged by adding 20 μM of RNA_15_ to 5 μM N. After 45 min, 3 μL was applied to glow-discharged Lacey grids (01895 – TedPella) and vitrified using a Vitrobot Mark IV system (Thermo Fisher Scientific) operated at 22 °C and 100% humidity. The images were acquired in a JEOL JEM 1400Plus 120 kV cryogenic transmission electron microscope equipped with a OneView camera 4k x 4k (Gatan).

### Zlotnick Analysis

#### (i) Determination of the nucleus size

Normalization was previously done using the results obtained from the sum of two exponentials with a linear drift. For this, first the linear drift was subtracted from the mass progress curves obtained by 370 nm light scattering. Then normalization was done considering the total amplitude of the curve. The condensate fraction was supposed to be proportional to the scattering signal, and the concentration of free protomers was estimated by:

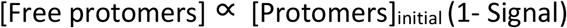

Plotting the log of the scattering signal (at *t*) vs the log of the free protomers (at *t*) and proceeding with a linear fitting yields the nucleus size from the slope. For RNA_15_ *t* was 30s and for tRNA 24s.

#### (ii) Determination of the elongation reaction order

Linear fittings were done for the linear region of the 370 nm light scattering traces. The log of these slopes was plotted against the log of the initial protomer concentration. The slope of these plot yields the elongation reaction order.

## Acknowledgments

We thank LNNano/CNPEM for access to the EM facility (Proposals 20233398 and 20231800). RVP and MAAA thank to FAPESP (Grants 2020/06062-1 and 2022/05088-2) We thank Chan Zuckerberg Initiative (CZI) for supporting RPM visit to LNNano/CNPEM (Grant 2021-240156/5022)

---

**Figure S1:**
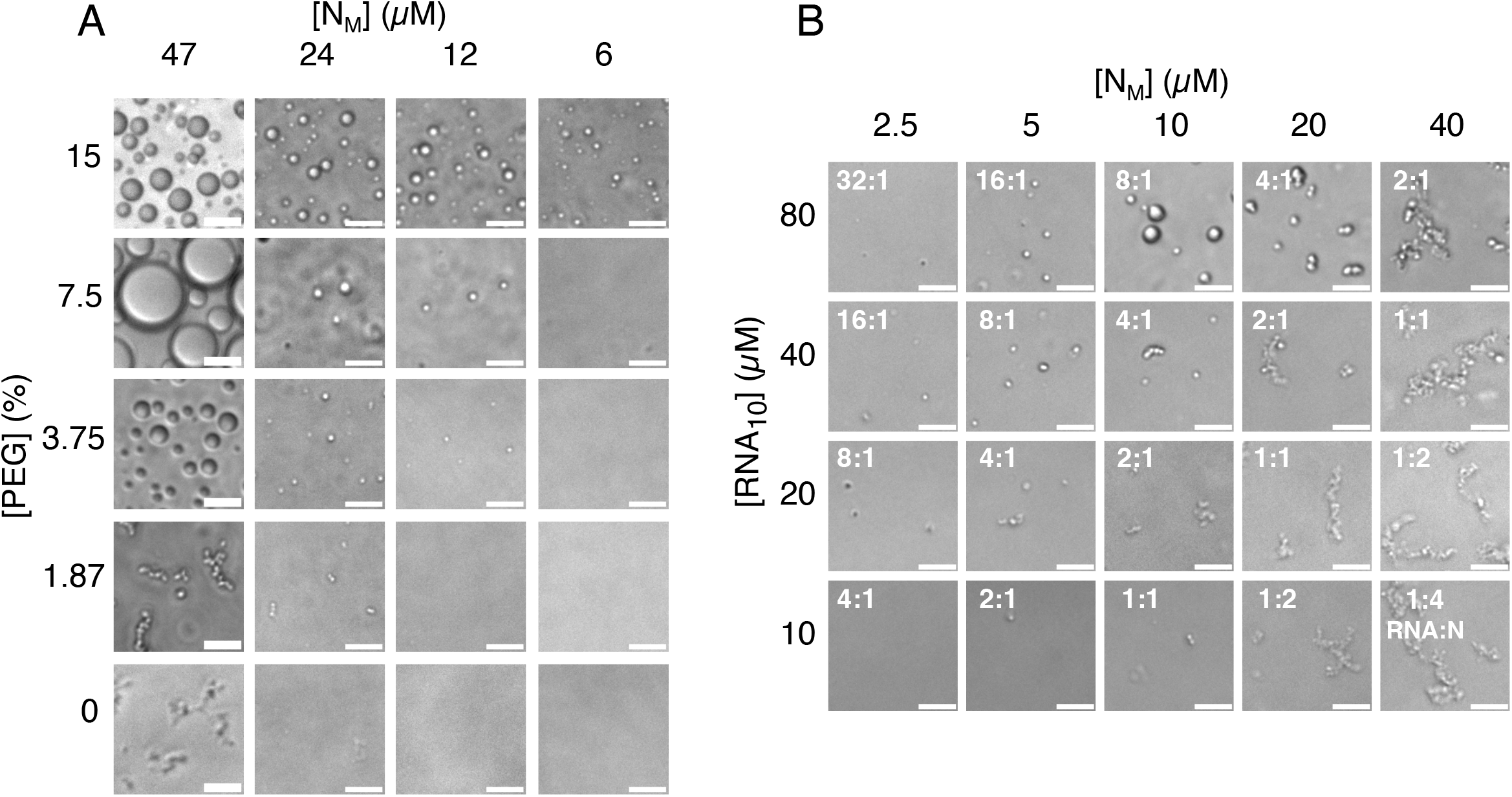
All protein concentrations used in this figure refer to monomer concentration [N_M_]. **(A)** Light microscopy images of homotypic condensates formed in presence of different concentrations of protein and crowding agent PEG 4000. At high protein concentrations (47 uM) when PEG is increased (7.5%) big coalesed condensates can be seen. Interestingly, prior to the formation of well rounded condensates (3.75% PEG) bran ched like con densates are formed (1.87% PEG), suggesting that a non homogeneous structure forms previous to the rounded condensates. **(B)** Light microscopy images showing heterotypic condensates formed in different concentrations of protein and RNA_10_.. For this, HEPES 20 mM pH 7.5, NaCl 50 mM was used, indicating that only marginal condensation can be seen with RNA_10_ and only in low ionic strength.

**Figure S2:**
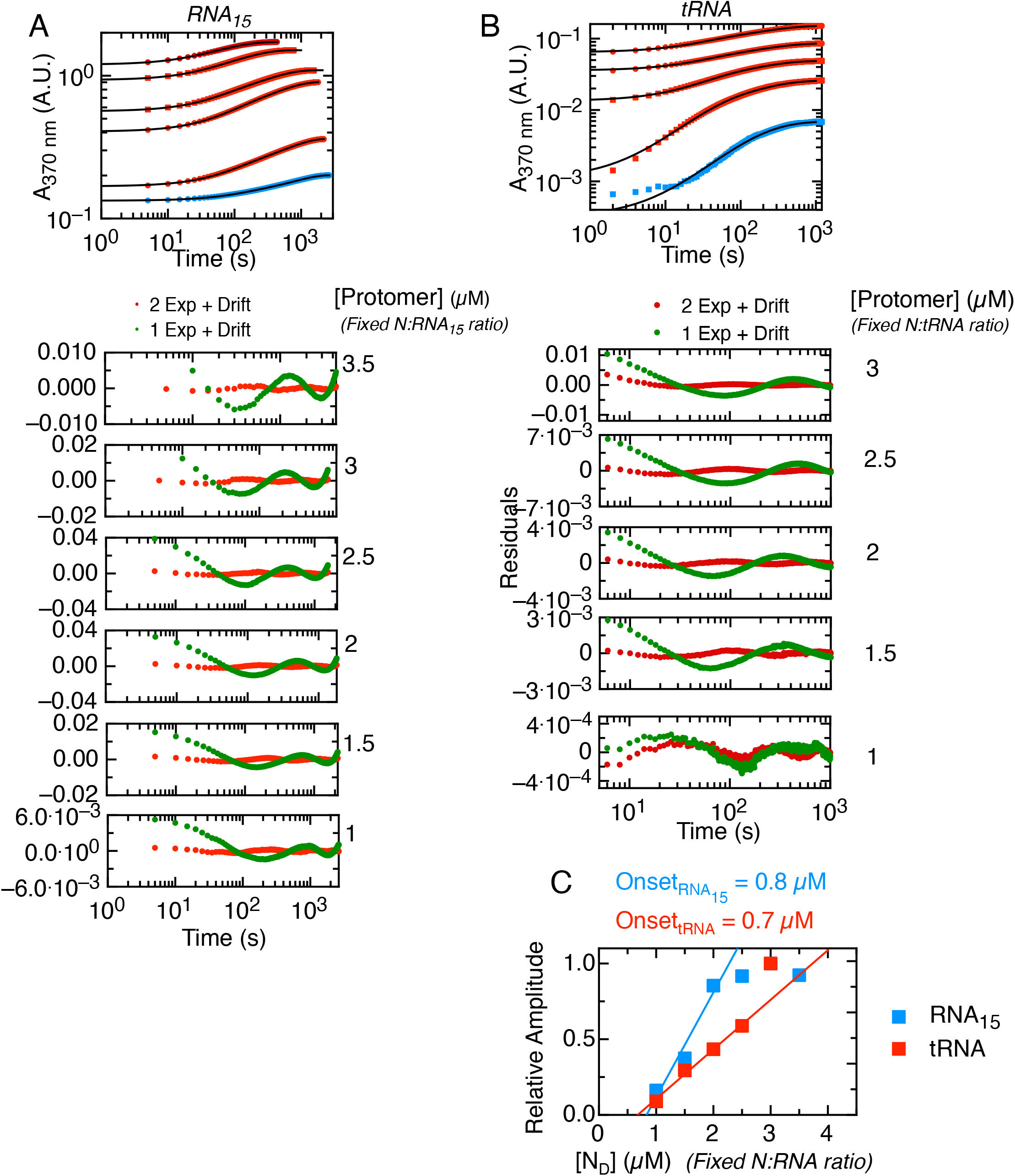
As model analysis depends on protein oligomeric state, all protein concentrations used in this figure refer to dimer concentration [N_D_]. Double exponential fittings of 370 nm scattering condensation traces. (C-D) Relative total amplitude of N-RNA_15_ and tRNA condensation traces shown in A and B. Interestingly, the onset concentration obtained fro the linear extrapolation to zero is in excellent agreement with the one obtained through Crystallization-like assembly model.

**Figure S3:**
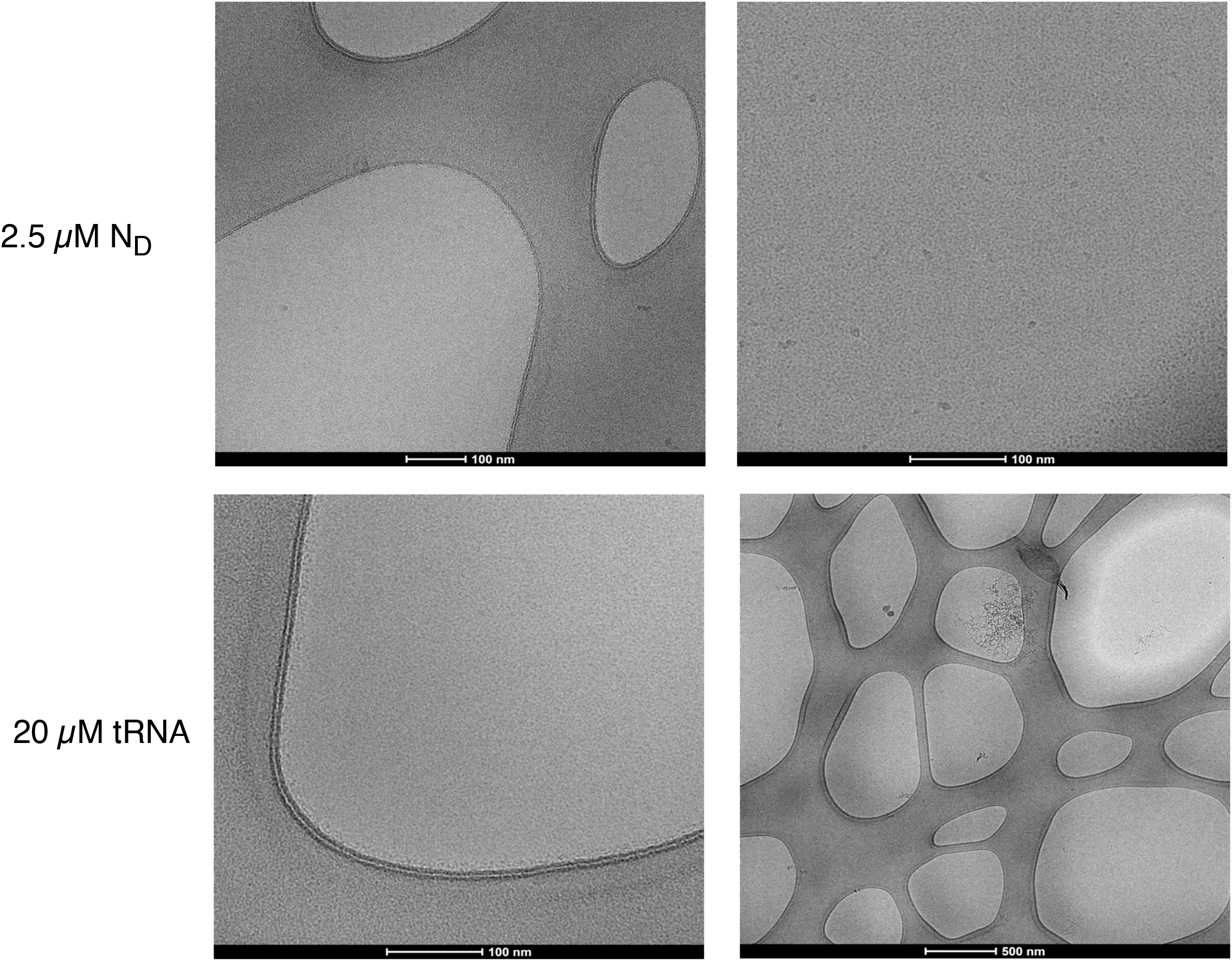
Cryo-EM controls. All protein concentrations used in this figure refer to dimer concentration [N_D_]. The holes (bubble shaped) seen in the images correspond to the Lacey grid holes. Neither the 5 uM N and 20 uM RNA_15_ yielded any visible structures.

